# Astrovirus replication is dependent on induction of double membrane vesicles through a PI3K-dependent, LC3-independent pathway

**DOI:** 10.1101/2023.04.11.536492

**Authors:** Theresa Bub, Virginia Hargest, Shaoyuan Tan, Maria Smith, Ana Vazquez-Pagan, Tim Flerlage, Pamela H. Brigleb, Victoria Meliopoulos, Brett Lindenbach, Valerie Cortez, Jeremy Chase Crawford, Stacey Schultz-Cherry

**Affiliations:** Department of Infectious Diseases, St. Jude Children’s Research Hospital, Memphis, Tennessee, USA; Integrated Program of Biomedical Sciences Department of Microbiology, Immunology, and Biochemistry, University of Tennessee Health Science Center, Memphis, Tennessee, USA; Graduate School of Biomedical Sciences, St. Jude Children’s Research Hospital, Memphis, Tennessee, USA; Department of Microbial Pathogenesis, Yale University, New Haven, Connecticut, USA; Department of Comparative Medicine, Yale University, New Haven, Connecticut, USA; Department of Molecular, Cellular and Developmental Biology, University of California, Santa Cruz, Santa Cruz, California, USA; Department of Immunology, St. Jude Children’s Research Hospital, Memphis, Tennessee, USA

**Keywords:** Astrovirus, autophagy, class III PI3K, double membrane vesicle, replication organelle

## Abstract

Human astrovirus is a positive sense, single stranded RNA virus. Astrovirus infection causes gastrointestinal symptoms and can lead to encephalitis in immunocompromised patients. Positive strand RNA viruses typically utilize host intracellular membranes to form replication organelles, which are potential antiviral targets. Many of these replication organelles are double membrane vesicles (DMVs). Here we show that astrovirus infection leads to an increase in DMV formation, and this process is replication-dependent. Our data suggest that astrovirus infection induces rearrangement of endoplasmic reticulum fragments, which may become the origin for DMV formation. Transcriptional data suggested that formation of DMVs during astrovirus infection requires some early components of the autophagy machinery. Results indicate that the upstream class III phosphatidylinositol 3-kinase (PI3K) complex, but not LC3 conjugation machinery, is utilized in DMV formation. Inhibition of the PI3K complex leads to significant reduction in viral replication and release from cells. Elucidating the role of autophagy machinery in DMV formation during astrovirus infection reveals a potential target for therapeutic intervention for immunocompromised patients.

**Importance:** These studies provide critical new evidence that astrovirus replication requires formation of double membrane vesicles, which utilize class III PI3K, but not LC3 conjugation autophagy machinery for biogenesis. These results are consistent with replication mechanisms for other positive sense RNA viruses. This suggests that targeting PI3K could be a promising therapeutic option for not only astrovirus, but other positive sense RNA virus infections.

## Introduction

Astroviruses are positive sense, single stranded, non-enveloped RNA viruses that cause disease in a variety of mammals and birds (1–4). In humans, infection is often associated with gastrointestinal symptoms such as nausea, vomiting, loss of appetite, stomach aches, and diarrhea (4–7). However, astrovirus infections can also result in fatal encephalitis, particularly in immunocompromised individuals (2, 7–12). Astrovirus infections are typically under-reported despite high prevalence (13, 14) Despite this, there are significant gaps in knowledge about astrovirus pathogenesis including the mechanisms behind viral replication.

Many positive sense, single-stranded RNA viruses including hepatitis C virus (HCV), coronaviruses, picornaviruses, and noroviruses utilize double membrane vesicles (DMVs) as replication chambers during infection. These replication organelles shield viral RNA from recognition by intracellular pattern recognition receptors (PRRs) that could alert the immune system (15–22). It has been suggested that these DMVs form with the aid of autophagy machinery. Generally, autophagy serves as the recycling system of the cell. During autophagy, double membrane vesicles called autophagosomes deliver cytoplasmic material to the lysosome for degradation. Autophagy can also be selective, targeting specific cargo such as depolarized mitochondria, damaged endoplasmic reticulum (ER) fragments, and others. The lysosome then fuses with the autophagosome, and cargo is degraded due to lysosomal enzymatic activity and acidic pH, recycling it for further use by the cell (23–26). The formation of the autophagosome can vary depending on whether the pathway is canonical or non-canonical. The canonical autophagy pathway involves machinery first characterized in starvation-induced autophagy, including the ULK1 pre-initiation complex, the class III Phosphatidylinositol 3-kinase (PI3K) complex required for production of phosphatidylinositol 3-phosphate (PI3P), and the LC3 conjugation system required for autophagosome maturation. Non-canonical pathways may utilize only some parts of the originally characterized autophagy machinery (27). Regardless of the pathway, these cellular components are often manipulated by viruses during infection to enhance viral replication.

Positive strand RNA viruses can hijack components of the autophagy machinery to form DMVs, which share characteristics with autophagosomes. However, viral-induced DMVs can be distinct from autophagosomes. They are not always delivered to lysosomes for degradation, tend to be smaller in size compared to autophagosomes, and importantly canonical autophagy machinery is not necessarily involved in the formation of these vesicles (15, 17–20, 22). Notably, viruses can induce the formation of DMVs from the ER, Golgi apparatus, mitochondria and other sites in the cell, which may have virus-specific implications for antiviral therapies (19, 22, 28).Recent evidence has shown that DMV formation is independent of LC3 lipidation machinery in both severe acute respiratory syndrome coronavirus 2 (SARS-CoV-2) and HCV infection; instead, both viruses appear to rely on the PI3K complex for formation of phosphatidylinositol 3-phosphate (PI3P) in DMV membranes (21, 29). Coxsackievirus B3 (CVB3), on the other hand, induces DMV formation that is independent of PI3K and ULK1 machinery, relying instead upon PI4KIIIβ for formation of these vesicles (30–32). Although electron microscopy images have shown association of lamb astrovirus with DMVs in lamb intestines (33), the role of autophagy in astrovirus infection has remained uncharacterized in both humans and animal models.

In the present study, we find that DMVs formed during astrovirus infection rely on a PI3K-dependent, LC3-independent autophagy pathway and may originate from the ER. This machinery is targetable, and using an autophagy specific PI3K inhibitor significantly reduces DMV formation and astrovirus replication. Targeting DMV formation through inhibition of the PI3K complex during astrovirus infection offers a potential therapy for astrovirus infection, and this therapy may further be applicable to other positive sense RNA viruses.

## Results

### Astrovirus induces DMV formation during replication

A previous study showed that astrovirus infection in lambs resulted in the formation of DMVs To determine if this was also true with human astroviruses, we performed transmission electron microscopy (TEM) on mock and human astrovirus-1 (HAstV-1) infected Caco-2 cells at 8-, 12-, 24-, and 36-hours post-infection (hpi). Beginning at 24 hpi, HAstV-1 infected cells had widespread formation of DMVs of approximately 200-500 nm in size compared to mock-inoculated Caco-2 cells (Fig. 1a). The DMVs were associated with HAstV-1 virions. Induction of DMV formation was dependent on productive viral replication, as UV-inactivated virus failed to induce DMVs (Fig. 1b).

**Fig 1.**
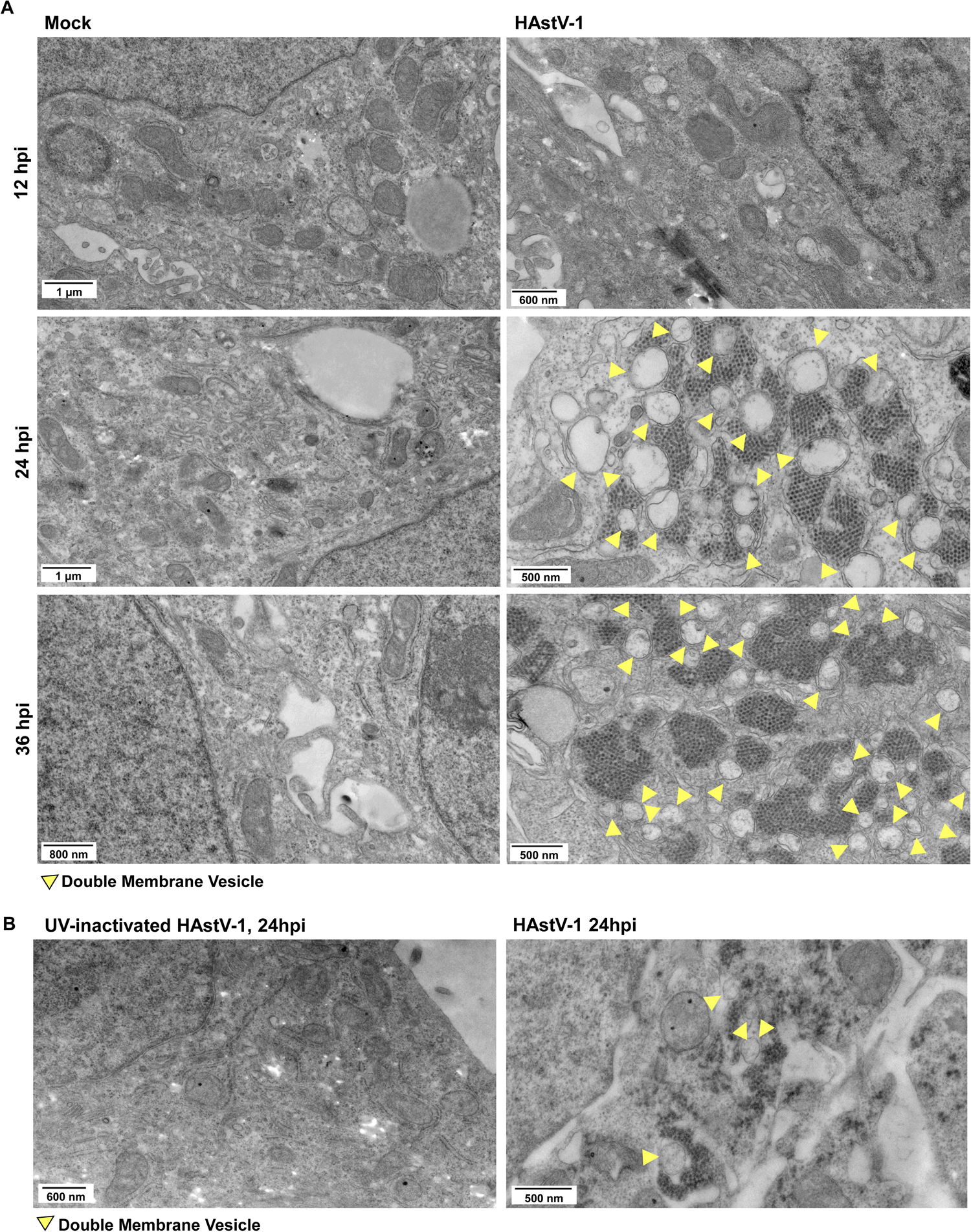
Astrovirus associates with DMVs during infection. (A) TEM images of mock-inoculated or HAstV-1-infected Caco-2 cells at 12, 24, and 36 hours post-infection (hpi). (B) TEM images of UV-inactive HAstV-1-inoculated Caco-2 cells and HAstV-1-infected Caco-2 cells at 24 hpi. (A & B) Yellow arrows indicate (DMVs).

### Astrovirus-induced DMVs may originate from the ER

To determine what cellular machinery is involved in DMV formation during astrovirus infection, we performed single cell RNA sequencing on astrovirus-infected Caco-2 cells at 4, 8, and 24 hpi (Fig. S1, Table 1). At 24 hpi, most cells in the HAstV-1-infected Caco-2 sample were infected and clustered together (Fig. 2a). The dataset also showed a significant upregulation in WHAMM expression at 24 hpi in HAstV-1 infected cells compared to other groups (Fig. 2b). WHAMM aids in actin nucleation and has been suggested to play a role in phagophore formation from the ER We found that WHAMM protein expression was also upregulated in HAstV-1 infected cells at 24 hpi compared to mock (Fig. 2c). Thus, we revisited our TEM images and identified multiple instances of ER fragments associated with DMVs during HAstV-1 infection in Caco-2 cells at 24 hpi (Fig. 2d). Confocal microscopy showed an association of the ER marker calnexin with viral dsRNA in HAstV-1-infected Caco-2 cells. In bystander cells, calnexin appeared to be more evenly distributed throughout the cytoplasm, whereas in infected cells, calnexin was clustered more closely around dsRNA, as evidenced by staining with the J2 antibody (Fig. 2e). This suggests that the ER could be a source of DMV membranes, which has been observed with other positive sense RNA viruses.

**Fig 2.**
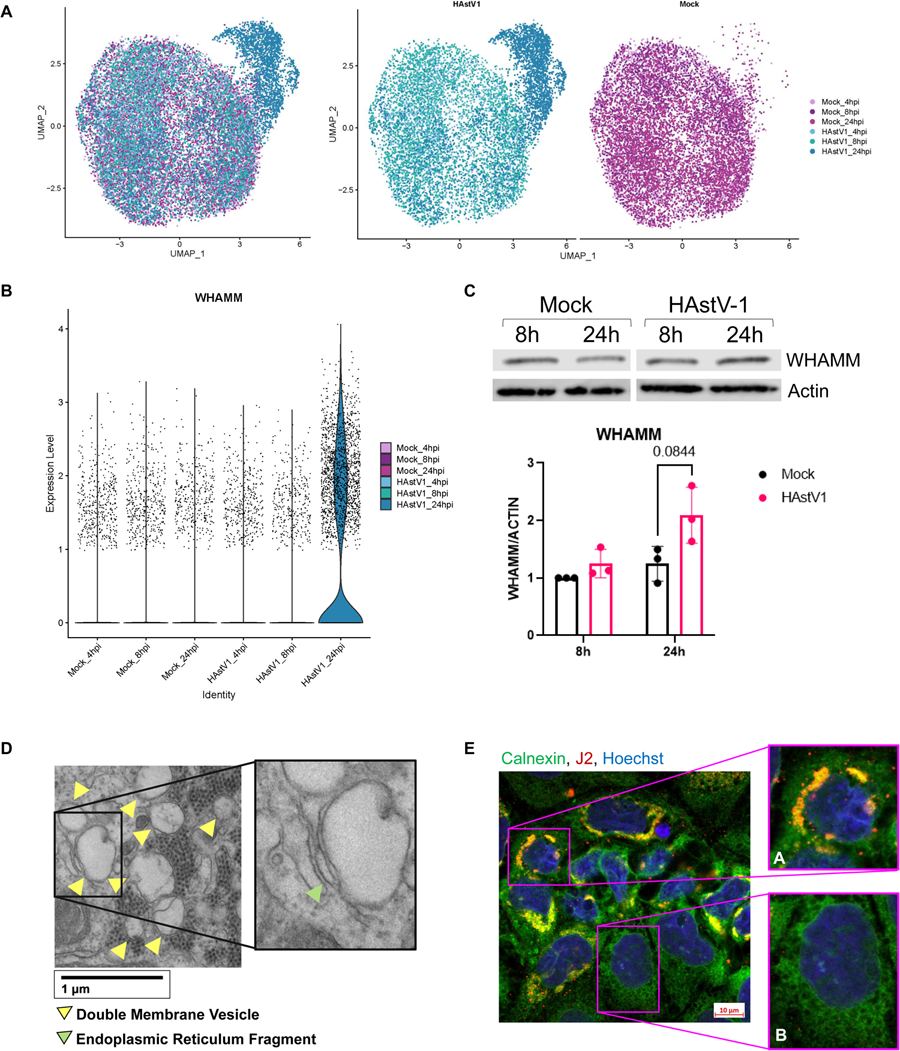
Single cell RNA sequencing shows dysregulation of autophagy-related genes in HAstV-1-infected cells at 24 hpi. (A) UMAP of 10X single cell RNA sequencing clusters by sample. (B) Violin plot showing expression level of WHAMM gene in each sample of the single cell RNA sequencing data set. (C) Immunoblot of WHAMM expression at 8 and 24 hpi in mock-inoculated and HAstV-1-infected Caco-2 cell lysates. Quantification of 8h and 24h time points is shown with statistical analysis by two-way ANOVA followed by Tukey’s multiple comparison test. (D) TEM image of HAstV-1-infected Caco-2 cells at 24 hpi. Yellow arrows indicate DMVs, and green arrow indicates ER fragment. (E) Confocal microscopy staining of HAstV-1-infected Caco-2 cells at 24 hpi, with Calnexin in green, dsRNA (J2) in red, and nucleus (Hoescht) in blue.

**Table 1:** Top 15 upregulated genes in each cluster for the HAstV-1 Caco-2 single cell RNA sequencing dataset

| Gene | cluster | avg_log2FC | pct.1 | pct.2 | p_val | p_val_adj |
| --- | --- | --- | --- | --- | --- | --- |
| KIAA1217 | 0 | 1.19850239 | 0.816 | 0.462 | 0 | 0 |
| LRP2 | 0 | 1.121160874 | 0.637 | 0.265 | 0 | 0 |
| CTNNB1 | 0 | 1.100106864 | 0.993 | 0.798 | 0 | 0 |
| CNN3 | 0 | 1.051849981 | 0.813 | 0.451 | 0 | 0 |
| APOBEC3C | 0 | 0.868743747 | 0.629 | 0.315 | 0 | 0 |
| PPP1R2 | 0 | 0.796207631 | 0.766 | 0.429 | 0 | 0 |
| CALR | 0 | 0.746712293 | 0.997 | 0.859 | 0 | 0 |
| SLC12A2 | 0 | 0.716473872 | 0.95 | 0.761 | 0 | 0 |
| ATP1B1 | 0 | 0.712313357 | 0.999 | 0.942 | 0 | 0 |
| TCF4 | 0 | 0.711609951 | 0.56 | 0.264 | 0 | 0 |
| AC106707.1 | 0 | 0.688501596 | 0.737 | 0.42 | 0 | 0 |
| FANCD2 | 0 | 0.75090556 | 0.572 | 0.286 | 9.19E-305 | 1.54E-300 |
| OLR1 | 0 | 0.814272779 | 0.403 | 0.171 | 3.81E-256 | 6.37E-252 |
| LINC00668 | 0 | 0.708536965 | 0.349 | 0.134 | 9.63E-252 | 1.61E-247 |
| CCND2 | 0 | 1.082390721 | 0.668 | 0.463 | 1.80E-247 | 3.02E-243 |
| GOLIM4 | 1 | 1.251522644 | 0.893 | 0.535 | 0 | 0 |
| HNF4A | 1 | 1.160206407 | 0.994 | 0.851 | 0 | 0 |
| PLAC8 | 1 | 1.119408282 | 0.549 | 0.203 | 0 | 0 |
| AGMAT | 1 | 1.015816816 | 0.743 | 0.399 | 0 | 0 |
| AFMID | 1 | 0.975713526 | 0.812 | 0.491 | 0 | 0 |
| RPS25 | 1 | 0.959045822 | 1 | 0.971 | 0 | 0 |
| SUMF2 | 1 | 0.909002623 | 0.895 | 0.563 | 0 | 0 |
| PHKG1 | 1 | 0.882683317 | 0.919 | 0.621 | 0 | 0 |
| **RPL23** | 1 | 0.858368645 | 1 | 0.982 | 0 | 0 |
| **CALR** | 1 | 0.857931888 | 0.998 | 0.873 | 0 | 0 |
| **SPTBN1** | 1 | 0.783309275 | 0.972 | 0.849 | 0 | 0 |
| **ALDH1B1** | 1 | 0.822800653 | 0.519 | 0.247 | 1.43E-263 | 2.40E-259 |
| **EGR1** | 1 | 0.779468435 | 0.811 | 0.583 | 2.59E-258 | 4.33E-254 |
| **INSR** | 1 | 0.76294615 | 0.742 | 0.462 | 1.44E-257 | 2.42E-253 |
| **SIPA1L3** | 1 | 0.765920406 | 0.597 | 0.317 | 2.69E-250 | 4.50E-246 |
| **KRT8** | 2 | 0.719943983 | 0.967 | 0.94 | 0 | 0 |
| **CD99** | 2 | 0.72737298 | 0.98 | 0.933 | 1.47E-283 | 2.47E-279 |
| **S100A14** | 2 | 1.218905825 | 0.813 | 0.689 | 3.15E-249 | 5.27E-245 |
| **GAPDH** | 2 | 0.690070593 | 0.874 | 0.833 | 1.52E-205 | 2.54E-201 |
| **DKK1** | 2 | 0.858903003 | 0.735 | 0.59 | 5.56E-194 | 9.30E-190 |
| **ITM2B** | 2 | 0.746170499 | 0.796 | 0.709 | 1.53E-193 | 2.56E-189 |
| **PRSS23** | 2 | 0.853212545 | 0.629 | 0.48 | 5.49E-164 | 9.20E-160 |
| **RRBP1** | 2 | 0.724677577 | 0.726 | 0.645 | 2.41E-139 | 4.04E-135 |
| **CADM1** | 2 | 0.743952903 | 0.643 | 0.53 | 6.48E-131 | 1.09E-126 |
| **BCAT1** | 2 | 0.67801791 | 0.717 | 0.622 | 2.73E-123 | 4.57E-119 |
| **H2AFZ** | 2 | 0.777433878 | 0.608 | 0.51 | 1.70E-117 | 2.85E-113 |
| **PKM** | 2 | 0.680011151 | 0.428 | 0.35 | 8.16E-60 | 1.37E-55 |
| **ZWINT** | 2 | 0.719273267 | 0.464 | 0.388 | 5.30E-59 | 8.87E-55 |
| **COTL1** | 2 | 0.680913965 | 0.426 | 0.349 | 5.93E-58 | 9.93E-54 |
| **BMP7** | 2 | 0.676959669 | 0.345 | 0.266 | 6.79E-50 | 1.14E-45 |
| **LGALS3** | 3 | 1.356751004 | 0.714 | 0.441 | 0 | 0 |
| **MT-CO1** | 3 | 0.596992212 | 0.991 | 0.979 | 4.74E-264 | 7.94E-260 |
| **RPL37** | 3 | 0.611049803 | 0.97 | 0.953 | 8.28E-216 | 1.39E-211 |
| **AGR2** | 3 | 0.858091 | 0.524 | 0.369 | 6.46E-124 | 1.08E-119 |
| **APOA1** | 3 | 1.112975286 | 0.411 | 0.265 | 1.59E-105 | 2.66E-101 |
| **CKB** | 3 | 0.779295318 | 0.656 | 0.582 | 7.28E-104 | 1.22E-99 |
| **RBP4** | 3 | 0.9377437 | 0.485 | 0.352 | 1.07E-102 | 1.80E-98 |
| **GOLIM4** | 3 | 0.620485939 | 0.681 | 0.593 | 9.25E-90 | 1.55E-85 |
| **VTN** | 3 | 0.922710756 | 0.354 | 0.226 | 1.83E-86 | 3.06E-82 |
| **TNFRSF11B** | 3 | 1.0122444 | 0.297 | 0.181 | 7.39E-72 | 1.24E-67 |
| **STARD10** | 3 | 0.608300202 | 0.618 | 0.561 | 3.93E-71 | 6.58E-67 |
| **SERPINA1** | 3 | 0.827951783 | 0.289 | 0.177 | 3.49E-67 | 5.84E-63 |
| **SLC26A2** | 3 | 0.744888023 | 0.3 | 0.209 | 2.48E-49 | 4.15E-45 |
| **ANXA4** | 3 | 0.694612524 | 0.257 | 0.18 | 1.15E-38 | 1.92E-34 |
| **FKBP11** | 3 | 0.580239191 | 0.257 | 0.205 | 8.98E-23 | 1.50E-18 |
| **HAstV1** | 4 | 3.733942054 | 0.898 | 0.184 | 0 | 0 |
| **ANKRD1** | 4 | 2.698331492 | 0.715 | 0.23 | 0 | 0 |
| **TFAP2C** | 4 | 2.553280214 | 0.792 | 0.302 | 0 | 0 |
| **KLF6** | 4 | 2.544868957 | 0.785 | 0.242 | 0 | 0 |
| **IFI6** | 4 | 2.328706584 | 0.736 | 0.293 | 0 | 0 |
| **NABP1** | 4 | 2.2978288 | 0.895 | 0.537 | 0 | 0 |
| **IRF1** | 4 | 2.271132526 | 0.58 | 0.05 | 0 | 0 |
| **RASGEF1B** | 4 | 2.250799675 | 0.792 | 0.343 | 0 | 0 |
| **STAT2** | 4 | 2.203833519 | 0.806 | 0.412 | 0 | 0 |
| **ZC3HAV1** | 4 | 2.086184961 | 0.796 | 0.311 | 0 | 0 |
| **SDC4** | 4 | 2.082058542 | 0.964 | 0.889 | 0 | 0 |
| **HELZ2** | 4 | 2.069541141 | 0.556 | 0.05 | 0 | 0 |
| **DTX3L** | 4 | 2.06726012 | 0.767 | 0.321 | 0 | 0 |
| **IFIT3** | 4 | 2.056124184 | 0.469 | 0.049 | 0 | 0 |
| **APOL2** | 4 | 2.049143309 | 0.552 | 0.146 | 0 | 0 |
| **HNRNPL** | 5 | 2.034851644 | 0.938 | 0.42 | 0 | 0 |
| **AC092069.1** | 5 | 1.892063036 | 0.983 | 0.669 | 0 | 0 |
| **MTA2** | 5 | 1.725865018 | 0.745 | 0.221 | 0 | 0 |
| **TMED2** | 5 | 1.596541391 | 0.916 | 0.446 | 0 | 0 |
| **RAB12** | 5 | 1.554177094 | 0.582 | 0.139 | 0 | 0 |
| **EIF5A** | 5 | 1.487754318 | 0.996 | 0.727 | 0 | 0 |
| **CDK2** | 5 | 1.30190575 | 0.483 | 0.098 | 0 | 0 |
| **RPS29** | 5 | 1.362759794 | 0.882 | 0.455 | 4.80E-282 | 8.04E-278 |
| **AMFR** | 5 | 1.31220832 | 0.859 | 0.408 | 7.52E-245 | 1.26E-240 |
| **GATC** | 5 | 1.294854977 | 0.708 | 0.273 | 1.76E-243 | 2.94E-239 |
| **PDIA4** | 5 | 1.235016341 | 0.877 | 0.488 | 2.80E-219 | 4.68E-215 |
| **NORAD** | 5 | 1.190846217 | 0.882 | 0.503 | 1.57E-215 | 2.62E-211 |
| **ERH** | 5 | 1.19704942 | 0.59 | 0.2 | 2.31E-208 | 3.87E-204 |
| **ANKRD1** | 5 | 1.40647639 | 0.589 | 0.263 | 5.52E-139 | 9.25E-135 |
| **IFI6** | 5 | 1.409000276 | 0.651 | 0.322 | 1.64E-132 | 2.75E-128 |
| **RPL12** | 6 | 1.285759387 | 0.848 | 0.844 | 4.10E-103 | 6.86E-99 |
| **RPS6** | 6 | 1.160863987 | 0.789 | 0.82 | 6.98E-67 | 1.17E-62 |
| **RPL13A** | 6 | 1.015001369 | 0.811 | 0.868 | 2.44E-61 | 4.09E-57 |
| **HMGB1** | 6 | 0.963484801 | 0.829 | 0.831 | 2.43E-53 | 4.07E-49 |
| **FTL** | 6 | 0.922028042 | 0.709 | 0.776 | 2.08E-33 | 3.49E-29 |
| **RPL21** | 6 | 1.303047417 | 0.612 | 0.661 | 1.60E-30 | 2.67E-26 |
| **YBX1** | 6 | 0.996963214 | 0.583 | 0.614 | 1.61E-23 | 2.70E-19 |
| **FTH1** | 6 | 1.082838047 | 0.543 | 0.587 | 1.78E-18 | 2.97E-14 |
| **RPS17** | 6 | 0.901755893 | 0.558 | 0.626 | 2.40E-16 | 4.01E-12 |
| **NSA2** | 6 | 1.060429197 | 0.405 | 0.406 | 2.44E-10 | 4.08E-06 |
| **RPL37A** | 6 | 0.925057149 | 0.522 | 0.611 | 2.99E-10 | 5.01E-06 |
| **RPL22** | 6 | 0.892429642 | 0.463 | 0.526 | 1.27E-08 | 0.000213251 |
| **YBX3** | 6 | 1.001063295 | 0.422 | 0.471 | 4.65E-07 | 0.00778686 |
| **DSTN** | 6 | 0.902448288 | 0.391 | 0.435 | 1.43E-05 | 0.238630221 |
| **ATP6V1G1** | 6 | 0.896333332 | 0.311 | 0.349 | 0.008948239 | 1 |
| **IFIT2** | 7 | 5.479733499 | 0.979 | 0.048 | 0 | 0 |
| **CCL5** | 7 | 5.405519828 | 0.741 | 0.009 | 0 | 0 |
| **IFIT3** | 7 | 4.375018516 | 0.965 | 0.086 | 0 | 0 |
| **HERC5** | 7 | 4.300257528 | 0.895 | 0.073 | 0 | 0 |
| **IFNL1** | 7 | 3.847226421 | 0.685 | 0.006 | 0 | 0 |
| **OTUD1** | 7 | 3.395153374 | 0.678 | 0.031 | 0 | 0 |
| **IFIH1** | 7 | 3.292621159 | 0.811 | 0.058 | 0 | 0 |
| **CXCL10** | 7 | 3.225257476 | 0.664 | 0.015 | 0 | 0 |
| **PLAAT2** | 7 | 3.178269385 | 0.35 | 0.009 | 0 | 0 |
| **AL158206.1** | 7 | 3.38718694 | 0.643 | 0.057 | 1.43E-204 | 2.40E-200 |
| **APOL2** | 7 | 3.88360581 | 0.951 | 0.182 | 2.56E-167 | 4.28E-163 |
| **MXD1** | 7 | 4.142727123 | 0.881 | 0.154 | 7.40E-163 | 1.24E-158 |
| **ZC3HAV1** | 7 | 5.304880867 | 0.993 | 0.357 | 1.03E-122 | 1.72E-118 |
| **PPP1R15A** | 7 | 3.367388945 | 0.713 | 0.128 | 4.90E-114 | 8.21E-110 |
| **SDC4** | 7 | 3.786158705 | 1 | 0.896 | 2.54E-83 | 4.25E-79 |

### PI3K machinery, but not LC3 conjugation machinery, is upregulated during astrovirus infection

Like autophagosomes, DMVs have a double membrane and can utilize components of the autophagy machinery during formation. Therefore, to determine if astrovirus-induced DMV formation was accompanied by an upregulation in autophagy machinery, we utilized an RT-PCR array (custom Qiagen RT^2^ Profiler) of canonical and alternative autophagy-related genes, as well as cell death-related genes, vesicular trafficking genes, and exosome-related genes. At 24 hpi, there was a significant upregulation in autophagy-related genes in HAstV-1 infected cells including *ULK1, AMBRA1, UVRAG*, *SQSTM1*, and *GABARAPL1,* in addition to *IDO1*. However, *MAP1LC3A, ATG5,* and *ATG7,* genes associated with LC3 conjugation machinery, were unchanged (Fig. 3a).

**Fig 3.**
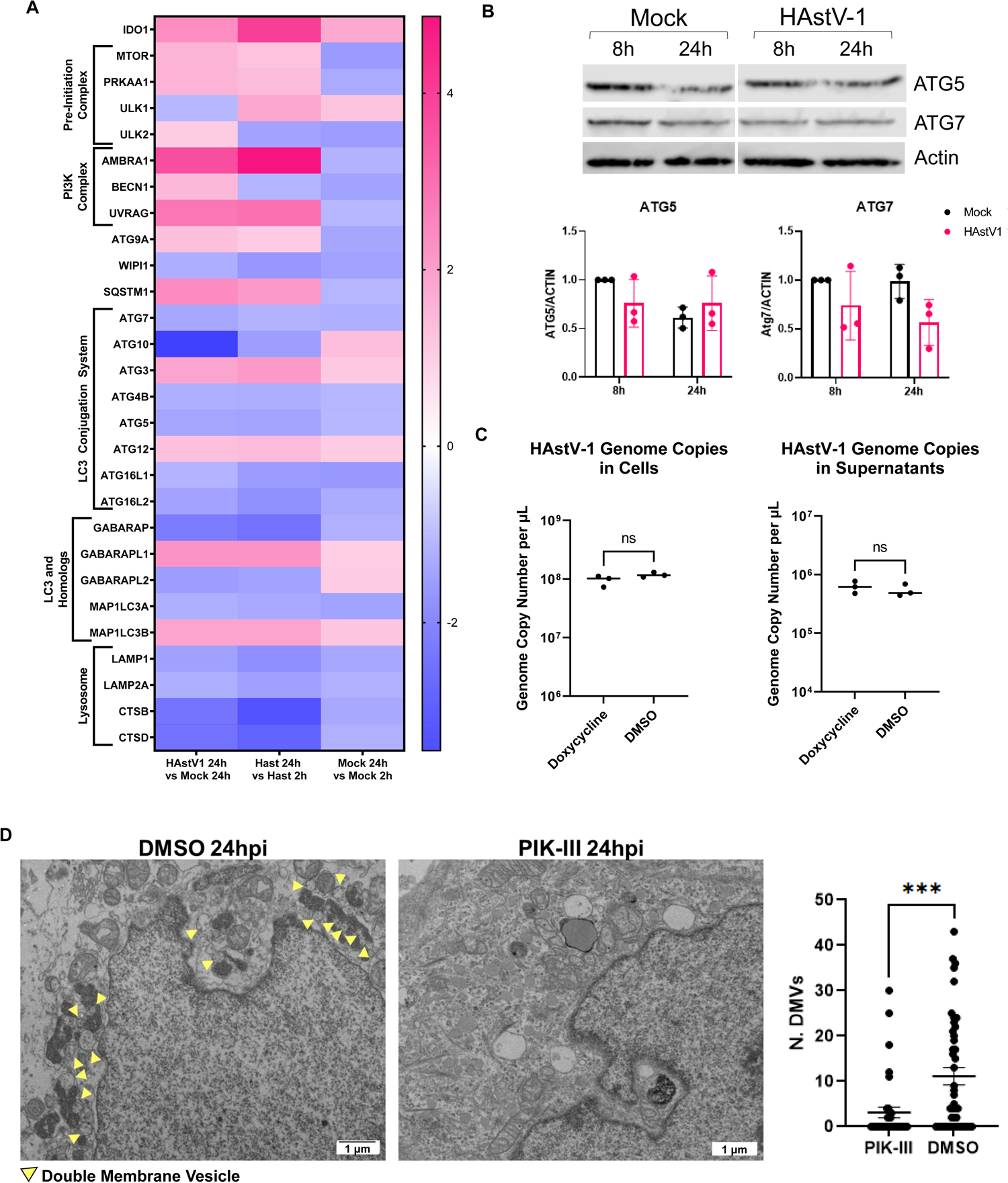
PI3K machinery, but not LC3 conjugation machinery, is upregulated during astrovirus infection. (A) Heat map of gene fold regulation from RT2 profiler dataset. (B) Immunoblot showing expression of ATG5 and ATG7 at 8 and 24 hpi in mock-inoculated and HAstV-1-infected Caco-2 cell lysates. Quantification of 8h and 24h time points was performed with statistical analysis by two-way ANOVA followed by Tukey’s multiple comparison test. (C) Genome copies per µL of astrovirus in cell lysate and supernatant RNA collected from HAstV-1-infected Huh-7.5 cells, where cells treated with doxycycline had induced RavZ protease activity. Statistical analysis was by unpaired two-tailed t test (D) TEM of PIK-III-treated and DMSO control HAstV-1-infected Caco-2 cells at 24 hpi and quantification of number of DMVs. Arrows indicate DMVs. Statistical analysis was by unpaired two-tailed t test. ***, P ≤ 0.001;

Immunoblots of lysates from mock and HAstV-1-infected Caco-2 cells confirmed that ATG5 and ATG7 were not upregulated (Fig. 3b). Given that we did not observe an upregulation of ATG5 and ATG7, we hypothesized that astrovirus-induced DMVs form independently of LC3 conjugation machinery. To test this, we utilized Huh-7.5 cells expressing doxycycline-inducible RavZ cysteine protease. RavZ has been shown to cleave LC3, impairing its ability to become conjugated to phosphatidylethanolamine, leading to a reduction in autophagosome formation (35, 36). After validating that induction of RavZ expression decreases LC3 levels, as shown by immunoblot (Fig. S2), we induced RavZ activity and infected the cells with HAstV-1. After infection with HAstV-1, we collected RNA from cells and supernatants for quantification of HAstV-1 genome copies at 24hpi. There was no change in genome copies in the absence of LC3 lipidation activity, suggesting that LC3 lipidation machinery and production of LC3-II are not required for astrovirus replication (Fig. 3c)

A recent study of SARS-CoV-2 and HCV replication demonstrated that the PI3K complex involved in formation of PI3P during autophagy is necessary to formation of DMVs during viral replication (21). One study showed that pan-PI3K inhibitors wortmannin and LY294002 were effective in reducing HAstV-1 infection (37, 38). However, this study did not address which PI3K complex is necessary for astrovirus infection or which part of the replication pathway is affected by inhibition. To test whether the autophagy-specific PI3K complex is required for astrovirus replication, we infected Caco-2 cells with HAstV-1 and treated Caco-2 cells with PIK-III, a specific PI3K complex inhibitor, or DMSO (Dimethyl Sulfoxide) at 1 hpi. At 24 hpi, the cells were fixed for TEM. PIK-III treatment significantly reduced the presence of DMVs and viral particles at 24 hpi, suggesting that the PI3K complex may support viral replication via formation of DMVs (Fig. 3d).

### Inhibition of the PI3K complex significantly reduces astrovirus replication

To determine whether inhibition of the PI3K complex affects astrovirus replication, we infected Caco-2 cells with HAstV-1 and treated cells with varying concentrations of PIK-III or DMSO at 2 hours pre-infection or 1 hpi. At 24 hpi, we fixed and stained the cells for HAstV-1 capsid or dsRNA. We observed significantly less capsid and dsRNA staining in the PIK-III-treated Caco-2 cells compared to the DMSO control at 24 hpi, and this difference was dose-dependent. Pre-infection treatment with PIK-III was not significantly different from post-infection treatment (Fig. 4a–b). We then repeated the experiment, collecting cells and supernatants at 24 hpi and extracted RNA to quantify HAstV-1 genome copies. Similarly, we found that PIK-III decreased HAstV-1 genome copies in both cell lysates and supernatant significantly in a dose-dependent manner (Fig. 4c). Finally, using supernatants from these cells, we found that cells that had been treated with PIK-III after HAstV-1 infection produced significantly less infectious virus than cells treated with DMSO (Fig. 4d). These experiments suggest that the PI3K complex aids in viral replication through formation of DMVs.

**Fig 4.**
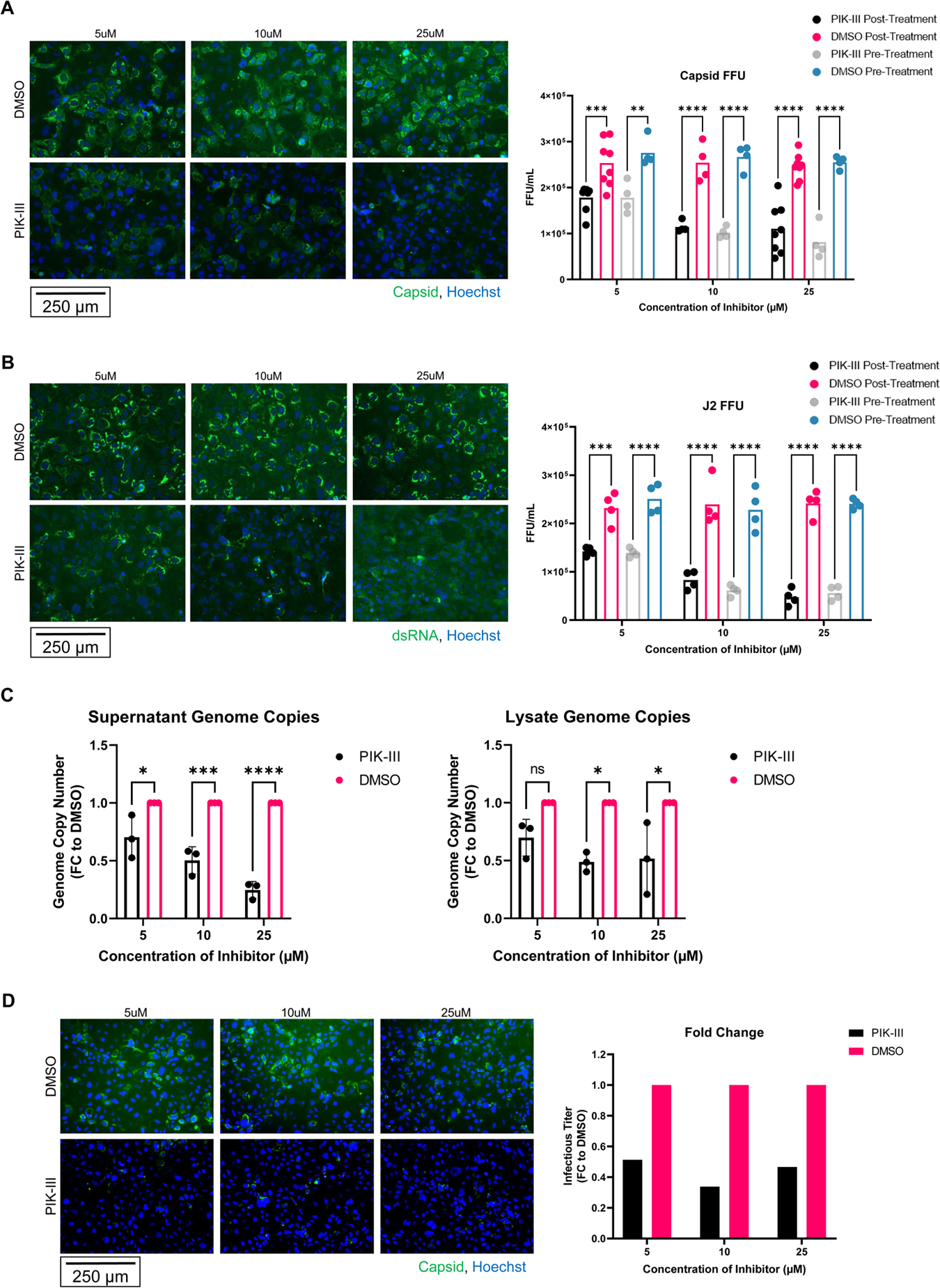
Inhibition of the PI3K complex reduces astrovirus replication. (A, B) HAstV-1-infected Caco-2 cells were either pre-treated for 2 hours prior to infection or treated 1 hpi with 5, 10, or 25 µM PIK-III or DMSO control (A) EVOS microscope images represent Caco-2 cells treated at 1 hpi, with astrovirus capsid in green and nucleus (Hoechst) in blue. Quantification shows FFU of samples treated 2 hours before infection and 1 hpi with statistical analysis by two-way ANOVA followed by Tukey’s multiple comparison test. (B) EVOS microscope images represent Caco-2 cells treated at 1 hpi, with astrovirus dsRNA (J2) in green and nucleus (Hoechst) in blue. Quantification shows FFU of samples treated 2 hours before infection and 1 hpi with statistical analysis by two-way ANOVA followed by Tukey’s multiple comparison test. (C) Genome copy number of human astrovirus in cell lysate and supernatant at 24 hpi from HAstV-1-infected Caco-2 cells treated with 5, 10, or 25 µM PIK-III or DMSO control at 1hpi with statistical analysis by two-way ANOVA followed by Tukey’s multiple comparison test. (D) Supernatants from HAstV-1-infected Caco-2 cells treated with 5, 10, or 25 µM PIK-III or DMSO control were collected and trypsin-treated at 24 hpi. Supernatants were used to infect Caco-2 cells. EVOS images show astrovirus capsid (green) and nucleus (Hoechst, blue). Quantification of FFU is shown. *, P ≤ 0.05; **, P ≤ 0.01; ***, P ≤ 0.001; ****, P ≤ 0.0001

### Single cell RNA sequencing suggests that dysregulation of autophagy machinery occurs only in astrovirus-infected cells, not bystander cells in vitro and in vivo

Using the single-cell RNA sequencing Caco-2 dataset, we confirmed that the upregulation of autophagy genes in HAstV-1-infected samples followed the same pattern as the RT-PCR array (Fig. 5a). In addition, upregulation in upstream autophagy pathway genes occurred only in astrovirus-infected but not bystander cells (Fig. 5b, S3). Using a single cell RNA sequencing dataset previously collected by our laboratory (13), we found that intestines from murine astrovirus-infected mice had an upregulation in *Pik3c3* in MuAstV-infected, but not bystander cells. This dataset showed a downregulation in *Map1lc3a* and its homologs *Gabarap* and *Gabarapl2*, as well as *Ulk1, Rb1cc1, Atg7, Atg10, Atg5,* and *Lamp1* in infected, but not bystander cells (Fig. 5c). This suggests that murine astrovirus replication could also utilize the PI3K complex, but not the LC3 conjugation machinery for replication *in vivo.* Future work will address the involvement of the PI3K complex in murine astrovirus replication and whether PI3K could be a therapeutic target for astrovirus infection spanning different species. These experiments provide evidence that astrovirus infection upregulates certain, but not all components of autophagy machinery *in vitro* and *in vivo* to facilitate viral replication (Fig. 5d).

**Fig 5.**
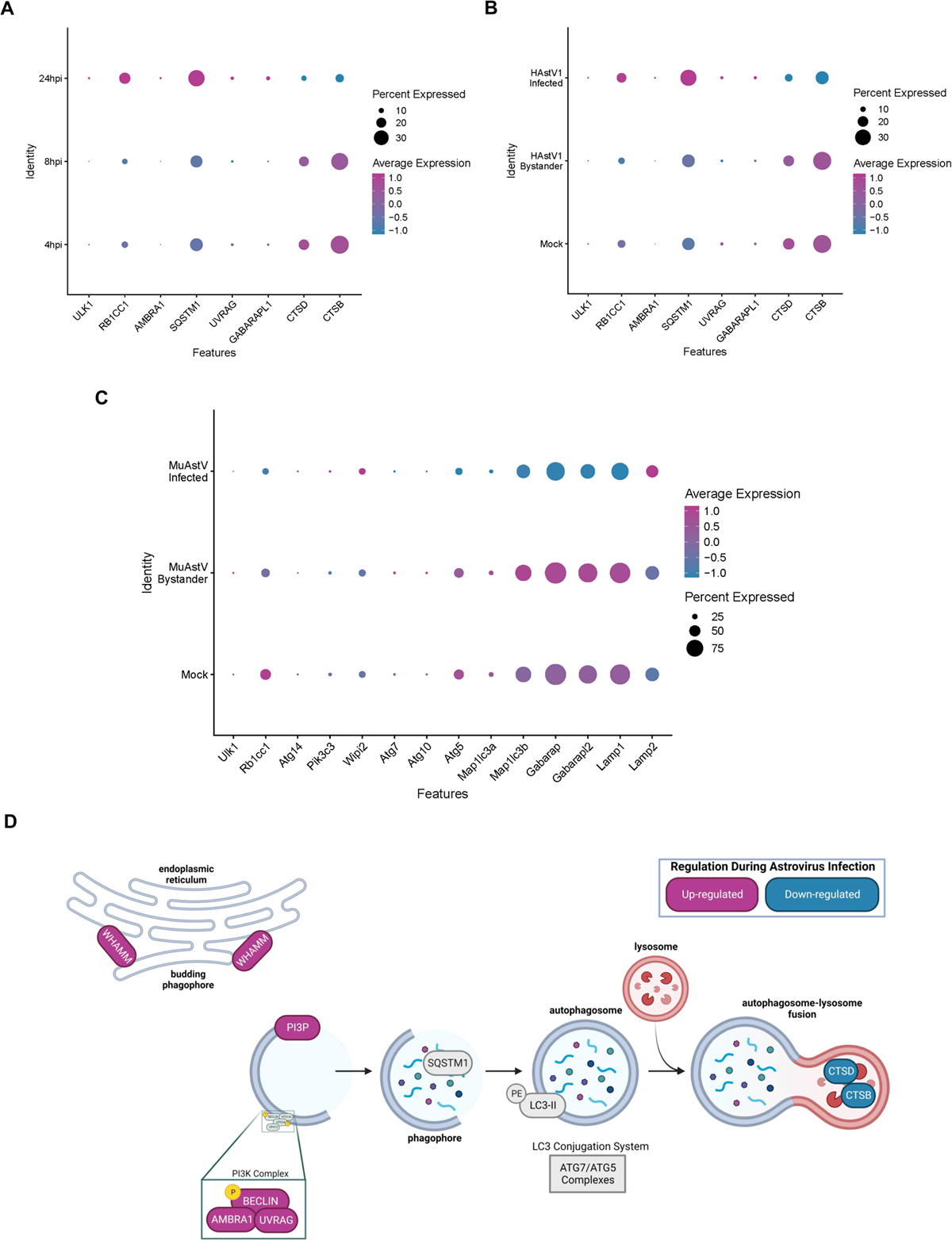
Single Cell RNA sequencing of HAstV-1-infected Caco-2 cells and MuAstV-infected mice shows dysregulation of autophagy genes. (A) Dot plot showing percent expression and average expression of autophagy-related genes in HAstV-1-infected Caco-2 cell samples at 4, 8, and 24 hpi from 10X single cell RNA sequencing dataset. (B) Dot plot showing percent expression and average expression of autophagy-related genes in HAstV-1-infected Caco-2 cells, HAstV-1 uninfected (bystander) Caco-2 cells, and mock-inoculated Caco-2 cells at 24 hpi from 10X single cell RNA sequencing dataset. (C) Dot plot showing percent expression and average expression of autophagy-related genes in MuAstV-infected, MuAstV uninfected (bystander), and mock-inoculated cells at 24 hpi from 10X single cell RNA sequencing murine astrovirus dataset (13). (D) Model of up-and down-regulated double membrane vesicle forming machinery during astrovirus infection.

## Discussion

Here, we show that astrovirus infection induces formation of DMVs, and this process is replication dependent. Formation of these DMVs also requires some, but not all canonical autophagy machinery. Previous studies of positive sense RNA virus replication have shown that canonical LC3 machinery may not be necessary for RNA virus replication using DMVs (21, 29). The LC3 conjugation system is indispensable for canonical, starvation-induced autophagy. It consists of *E1* and *E3*-like proteins ATG5 and ATG7, which work together to conjugate LC3-I to phosphatidylethanolamine (PE) to form LC3-II. This crucial step leads to autophagosome maturation (27, 39). Our work is consistent with previous literature, as we also observed that inhibition of LC3 machinery does not affect astrovirus replication.

In addition to validating these previous findings, we furthermore demonstrate that astrovirus uses the autophagy specific PI3K complex for formation of DMVs. The PI3K complex initiates phagophore formation and production of PI3P during canonical autophagy (27, 39). Inhibiting this complex greatly reduces astrovirus replication and infectious virus production, as well as formation of DMVs as seen by electron microscopy. These data are consistent with recent studies showing that SARS-CoV-2 and HCV utilize the PI3K complex, but not LC3 conjugation machinery for DMV formation and replication (21, 29). Our work suggests that early parts of autophagy machinery are essential for DMV formation, while LC3 conjugation is not required.

Without LC3 involvement in the formation of DMVs, it is possible that an LC3 homolog, such as the significantly upregulated GABARAPL1, could be active in the formation of DMVs during astrovirus replication. Notably, while inhibition of PI3K significantly reduces astrovirus replication, it does not ablate replication entirely. One possible explanation for this is that other phosphatidylinositol kinases are also involved in formation of these DMV replication organelles, such as PI4K (19, 21, 22, 24). Use of a PI4K inhibitor during astrovirus infection could determine whether this is the case.

While it is clear that formation of DMVs during astrovirus infection is replication-dependent, it is not yet determined which parts of the astrovirus genome are necessary for inducing formation of DMVs. Nonstructural proteins alone from other viruses such as HCV and SARS-CoV-2 are sufficient for induction of DMV formation (40–42). Little is known about the function of astrovirus nonstructural proteins, and it is likely that they play a role in DMV formation (43, 44).

While the formation of DMVs can occur using a variety of host cellular membranes, we have shown that ER-associated WHAMM is significantly upregulated during astrovirus infection. Future work will address whether WHAMM is necessary for budding of DMVs from ER membranes during astrovirus replication, or if other host membranes such as the Golgi will compensate in the absence of WHAMM. In addition, there are many different activators and inhibitors of canonical and non-canonical autophagy pathways, depending on different stressors activating the pathway such as starvation, genotoxic stress, and ER stress (45–49). Future work should also explore the role of mTOR, AMPK, and other regulators upstream of the autophagy pathway to determine whether these also affect DMV formation during astrovirus infection.

Altogether, our results indicate that astrovirus replication relies upon the formation of DMVs using early autophagy machinery, including the PI3K complex, but not LC3 conjugation machinery. Future studies will address whether the PI3K complex is necessary for astrovirus infection in brain cells, leading to encephalitis in immunocompromised populations. Astrovirus-induced DMVs appear to bud from the ER using the nucleation-promoting factor WHAMM. These results emphasize how distinct positive strand RNA viruses utilize similar mechanisms of replication. Although SARS-CoV-2, HCV, and HAstV-1 are different, their common use of the PI3K machinery implies the possibility of a conserved therapeutic target for many positive sense RNA viruses. Understanding these replication mechanisms will help to determine future antiviral therapies.

## Materials and Methods

### Cells and Virus Propagation

Caco-2 human intestinal adenocarcinoma cell line was obtained from ATCC (HTB-37). Cells were grown in Corning minimum essential medium (MEM) containing 20% fetal bovine serum (FBS; HyClone), GlutaMax (Gibco), 1mM sodium pyruvate (Gibco), and penicillin-streptomycin (Fisher).

The Huh-7.5 Rav-Z inducible cell line was a generous gift from Brett Lindenbach’s lab at the Yale School of Medicine. These cells were grown in DMEM (Thermo Fisher) containing 10% FBS (HyClone) and 3 µg/mL puromycin (Invitrogen).

HAstV-1 lab adapted viral stock was propagated in Caco-2 cells. Viral titer was quantified using fluorescent-focus assay (FFU) as previously described (50). For UV inactivation experiments, a UV cross-linker was utilized to subject HAstV-1 to 100 mJ/cm^2^, and inactivation was confirmed using FFU assay.

### Transmission Electron Microscopy

Caco-2 cells were plated in a 6-well plate (3.5 × 10^5^). After 46 hours, appropriate samples were treated with 10 µM PIK-III or DMSO. At 48 hours post-plating, cells were inoculated with supernatants taken from HAstV-1 (MOI 10) or mock-inoculated Caco-2 cells in serum free media for 1 hour. Following virus adsportion, inoculum was replaced with either fresh media, media containing 10 µM PIK-III, or media containing DMSO. At 8, 12, 24, or 36 hpi, cells were fixed in 2.5% Glutaraldehyde/2% PFA in 0.1M Cacodylate Buffer. Following fixation, samples were post fixed in osmium tetroxide and contrasted with aqueous uranyl acetate. Samples were dehydrated by an ascending series of ethanol to 100% followed by 100% propylene oxide. Samples were infiltrated with EmBed-812 and polymerized at 60°C. Embedded samples were sectioned at ∼70nm on a Leica (Wetzlar, Germany) ultramicrotome and examined in a ThermoFisher Scientific (Hillsboro, OR) TF20 transmission electron microscope at 80kV. Digital micrographs were captured with an Advanced Microscopy Techniques (Woburn, MA, USA) imaging system. Unless otherwise indicated, all reagents from Electron Microscopy Sciences (Hatfield, PA, USA).

### RT^2^ Profiler

Caco-2 cells were plated in a 6-well plate (3.5 × 10^5^). After 48 hours, cells were inoculated with supernatants taken from HAstV-1 (MOI 10) or mock-inoculated Caco-2 cells in serum free media for 1 hour. Following virus adsportion, the inoculum was replaced with fresh media. At 2, 8, and 24 hpi, cell supernatants were collected in TRIzol LS. Cells were collected in TRIzol, and RNA was extracted from all samples per manufacturer’s instructions. RNA quality was checked using Thermo Scientific NanoDrop 2000 per manufacturer’s instructions. Then qRT-PCR was performed on supernatant RNA to determine genome copies of HAstV-1 in supernatants, as previously described (51). We confirmed that genome copies of astrovirus increased in HAstV-1-infected cell supernatants over time, and no genome copies were detected in mock-inoculated cell supernatants. Then, RNA from cells was reverse transcribed using the RT^2^ First Strand Kit from Qiagen (Qiagen 330401). After cDNA was collected, it was utilized with SYBR Green qPCR Mastermix (Qiagen 330500) in a custom-designed real-time RT^2^ Profiler PCR Array (Qiagen 330171). CT values were collected, and data analysis was performed using Qiagen’s data analysis web portal (http://www.qiagen.com/geneglobe). In addition, we have found that astrovirus induces epithelial to mesenchymal transition, reducing epithelial markers on Caco-2 cells later in infection (52). Thus, *IDO1* and *EpCAM* were included in the panel to verify normal cellular response to astrovirus infection.

Mock and HAstV-1-treated samples were designated as control and test groups respectively. All samples passed quality checks. Reference genes were included in the RT^2^ panel, and data was normalized to these genes. The Qiagen data analysis protocol included fold change/regulation calculations based on ΔΔCT calculations. Statistical analysis on the Qiagen web portal utilized a student’s t-test to calculate p values, where parametric, unpaired, two-sample equal variance, two-tailed distribution was utilized.

### 10X Single Cell RNA Sequencing Sample Preparation

Caco-2 cells were plated in a 6-well plate (3.5 × 10^5^). Samples were assigned to wells corresponding to 4-, 8-, or 24-hours post-infection (hpi). At 48 hours post-plating, 24 hpi cell wells were inoculated with supernatants taken from HAstV-1 (MOI 10) or mock-inoculated Caco-2 cells in serum free media for 1 hour. After virus adsorption, the inoculum was replaced with fresh media. At 64 hours post-plating, 8 hpi cell wells were inoculated with supernatants taken from HAstV-1 (MOI 10) or mock-inoculated Caco-2 cells in serum free media for 1 hour. After virus adsorption, the inoculum was replaced with fresh media. At 68 hours post-plating, 4 hpi cell wells were inoculated with supernatants taken from HAstV-1 (MOI 10) or mock-inoculated Caco-2 cells in serum free media for 1 hour. After virus adsorption, the inoculum was replaced with fresh media. At 72 hours post-plating, all cells were washed with PBS and harvested using trypsin. Cells were filtered through at 70-µm cell strainer. Cells were spun down at 1200 rpm for 10 minutes at 4°C. Cells were washed in cell wash buffer (1% BSA in PBS) and spun down again at 1200 rpm for 10 minutes at 4°C. Finally, cells were resuspended in 50 µL of cell wash buffer and counted using a hemocytometer. Cells were resuspended in appropriate volume to reach 1000 cells/µL. Next, 9,000 cells were loaded onto the 10X Genomics Chromium controller for partitioning of single cells into gel beads with a goal of recovering 6,000 cells. Next, using a 10X Genomics 3’ Gene Expression Kit (version 3.1) according to manufacturer’s instructions, single-cell transcriptomics libraries were produced. Libraries were sequenced using Illumina NovaSeq 2000 at suggested sequencing lengths and depths.

### 10X Single Cell RNA Sequencing Analysis

The 10X transcriptomics data were first processed using CellRanger count (version6.1.1, 10X Genomics). GRCh38 was used as our reference, which was altered to include the human astrovirus 1 genome (accession ID MK059949.1). Samples were aggregated and normalized by the median number of mapped reads per identified cell using CellRanger aggr. Normalized gene-expression matrices were then imported into Seurat (version4.1.1) for downstream analysis and data visualization.

Data were first filtered by excluding any gene that was not present in at least 0.1% of total called cells (23 cells). Cells that exhibited extremes in the total number of transcripts expressed (>6000), the total number of genes expressed (<400 or >3000), or mitochondrial gene expression (>8%) were then excluded from downstream analyses. Data were log-normalized using default parameters. We identified the top 2,000 variable features using the vst method after excluding the astrovirus gene.

The fastMNN algorithm was then utilized to integrate datasets from distinct libraries, effectively minimizing subject-and sample-specific differences in order to identify similar transcriptional subsets. The first 25 fastMNN dimensions were used for UMAP dimensionality reduction and for nearest-neighbor graph construction for identifying transcriptional clusters in Seurat. Markers for each cluster were identified using FindAllMarkers function (min.pct = 0.25, logfc.threshold = 0.25) Differential gene expression was assessed using the default FindMarker function (min.pct = 0.01, logfc.threshold = 0.01). We generated a subset for 24 hpi samples alone. This subset was processed as described above.

### Immunofluorescent Staining

Caco-2 cells were plated on ibidi µ-Slide 8 Well high Polymer chamber slides at a density of 4 × 10^4^ cells per well. At 48 hours post-plating, cells were inoculated with supernatants taken from HAstV-1 (MOI 10) or mock-inoculated Caco-2 cells in serum free media for 1 hour. Following virus adsportion, inoculum was replaced with fresh media. At 24 hpi, cells were fixed in 4% paraformaldehyde (PFA) in PBS at room temperature for 20 minutes. Cells were washed in PBS. Next, 0.1% Triton X in PBS was utilized to permeabilize the cells for 15 minutes at room temperature. Cells were washed in PBS and blocked in 5% normal goat serum (NGS) in PBS for 1 hour at room temperature. Cells were washed in PBS, and antibodies were diluted in 1% NGS/PBS. Primary antibodies included Calnexin (Invitrogen PA5-34754) at 1:1000 and J2 (SciCons 10010500) at 1:100. Cells were incubated in antibody solution overnight at 4°C. The next day, samples were washed in PBS. Cells were incubated in secondary antibody solution for 45 minutes in the dark. Secondary antibodies included Alexa Fluor 488 goat anti-Rabbit (Invitrogen A11008) at 1:1000, Alexa Fluor 555 goat anti-Mouse (Invitrogen A21422) at 1:1000, and Hoechst (ThermoFisher H3569), 1:2000 in 1% NGS/PBS. Cells were again washed in PBS, and samples were finally fixed in Prolong Gold Antifade Mountant (Invitrogen). Prepared samples were imaged on a Zeiss LSM 780 Observer.Z1 using a Plan Apochromat 63X/1.4 objective lens. A 1024 × 1024 pixel array, final pixel size of 88nm, and pixel dwell time of 1.27 µs was used. Based on the green channel, a 1 AU pinhole size was selected. For each channel, gain was set to 500. A 405nm diode laser was utilized for Hoechst fluorescence, and 410-495 nm light was detected using an alkali PMT. A 488 nm multi-line Argon laser was utilized for Alexa Fluor 488 fluorescence, and 499-579 nm light was detected using a GaAsP PMT. Finally, a 561 nm DPSS laser was utilized for Alexa Fluor 555 fluorescence, and 588-712 nm light was detected using an alkali PMT. Acquisition was completed using Zen Black 2012 SP 5 (14.0.28.201).

For PIK-III experiments, Caco-2 cells were plated in a 96-well plate (2.5 × 10^4^). At 46 hours post-plating, appropriate wells were treated with 5, 10, or 25 µM PIK-III or DMSO control. At 48 hours post-plating, cells were inoculated with supernatants taken from HAstV-1 (MOI 10) or mock-inoculated Caco-2 cells in serum free media for 1 hour. Following virus adsportion, the inoculum was replaced with fresh media or media containing 5, 10, or 25 µM PIK-III or DMSO control. At 24 hpi, cells were fixed in 100% methanol. Cells were washed in PBS and incubated in primary antibody solution containing astrovirus capsid monoclonal antibody 8e7 (Invitrogen MA5-16293) at 1:100 in 1% NGS/PBS for 1 hour at room temperature. Cells were washed in PBS. Cells were incubated in secondary antibody solution containing Alexa Fluor 488 goat anti-Mouse (Invitrogen A10680) at 1:1000 and and Hoechst (ThermoFisher H3569) at 1:2000 in 1% NGS/PBS for 45 minutes in the dark. Cells were again washed in PBS. Samples were imaged using the EVOS FL cell imaging system and analyzed using ImageJ 2.9.0/1.53t software. FFU was calculated as previously described (S. Marvin et al., 2014). The same method was used for supernatants from PIK-III and DMSO-treated cell supernatants used to infect fresh Caco-2 cells in a 96-well plate.

### Immunoblotting

Caco-2 cells were plated in a 6-well plate (3.5 × 10^5^). At 48 hours post-plating, cells were inoculated with supernatants taken from HAstV-1 (MOI 10) or mock-inoculated Caco-2 cells in serum free media for 1 hour. Following virus adsportion, inoculum was replaced with fresh media. At the proper time point, 8, or 24 hpi, cells were collected in 250 µL RIPA Lysis Buffer (Abcam) containing 1x protease inhibitor cocktail (Pierce) on ice. Samples were vortexed briefly and kept on ice for 30 minutes. Samples were frozen at –80°C until used. Sample protein concentration was determined using BCA Protein Assay Kit (Pierce). Equal concentrations of protein were prepared under reducing conditions and separated by sodium dodecyl sulfate-polyacrylamide gel electrophoresis (SDS-PAGE) (4–20% Tris Glycine 1.0mm Mini Protein Gels from Invitrogen XP04200BOX). Gels were transferred to PVDF membranes using the iBlot™ 2 transfer stacks (ThermoFisher IB24002). Membranes were probed for protein with respective primary antibodies and IRDye 680RD goat anti-rabbit IgG secondary antibody using the ThermoFisher iBind device according to manufacturer’s instructions. Primary antibodies included β-Actin (Cell Signaling 4970S) at 1:1000, Atg5 (Abcam ab108327) at 1:1000, Atg7 (Abcam ab52472) at 1:1000, and WHAMM (Abcam ab122572) at 1:1000.

For Huh-7.5 immunoblots, Huh-7.5 cells were plated in a 6-well plate (3 × 10^5^). After 48 hours, cells were treated with 1.5 or 6 µM doxycycline to induce Rav-Z protease activity or DMSO controls. At 24 hours post-treatment, cells were inoculated with supernatants taken from HAstV-1-infected (MOI 10) Caco-2 cells in serum free media for 1 hour. Following virus adsorption, the inoculum was replaced with media containing the appropriate concentration of doxycycline or DMSO, as well as 30 µM Chloroquine to enable monitoring of LC3-II/LC3-I (53). At 24 hpi, cells were collected in 250 µL RIPA Lysis Buffer (Abcam) containing 1x protease inhibitor cocktail (Pierce) on ice. Samples were vortexed briefly and kept on ice for 30 minutes. Samples were frozen at –80°C until used. Sample protein concentration was determined using BCA Protein Assay Kit (Pierce). Immunoblots were performed the same way as described with Caco-2 cell lysates. Membranes were probed for LC3 (Cell Signaling 2775S) at 1:1000 overnight at 4°C in 5% BSA/TBST. The following day, membranes were incubated in secondary antibody solution containing IRDye 680RD goat anti-rabbit IgG secondary antibody in 5% BSA/TBST for 1 hour at room temperature. Membranes were imaged on the LI-COR Odyssey Fc (Software version number 1.0.36). Next, membranes were stained for β-Actin (Cell Signaling 4970S) at 1:1000 using the ThermoFisher iBind device as described above. All immunoblots were performed in triplicate. Each blot was imaged on the LI-COR Odyssey Fc. Densitometry was measured using Image Studio version 5.2 software.

### PIK-III and Astrovirus Infection RT-PCR

For PIK-III experiments, Caco-2 cells were plated in a 96-well plate (2.5 × 10^4^). At 48 hours post-plating, cells were inoculated with supernatants taken from HAstV-1 (MOI 10) or mock-inoculated Caco-2 cells in serum free media for 1 hour. Following virus adsportion, the inoculum was replaced with fresh media or media containing 5, 10, or 25 µM PIK-III or DMSO control. At 24 hpi, supernatants were collected in TRIzol LS, and cells were collected in TRIzol, and RNA was extracted per manufacturer’s instructions. RNA was then utilized in RT-PCR to determine astrovirus genome copies, as previously described (51).

### Huh-7.5 Rav-Z Induction

Huh-7.5 cells were plated in a 6-well plate (3 × 10^5^). After 48 hours, cells were treated with 3 µM doxycycline to induce Rav-Z protease activity or DMSO control. At 24 hours post-treatment, cells were inoculated with supernatants taken from HAstV-1-infected (MOI 10) Caco-2 cells in serum free media for 1 hour. Following virus adsorption, the inoculum was replaced with media containing the appropriate concentration of doxycycline or DMSO. At 24 hpi, supernatants were collected in TRIzol LS, and cells were collected in TRIzol. RNA was extracted from these samples according to manufacturer’s instructions. RNA was then utilized in RT-PCR to determine astrovirus genome copies, as previously described (51).

### MuAstV 10X Dataset

The murine astrovirus 10X single cell RNA sequencing dataset was collected previously, as described (13).

### Statistical Analysis

Data were analyzed by two-way ANOVA followed by Tukey’s multiple comparisons test (western blots, PIK-III genome copies, and PIK-III FFU analysis) or unpaired two-tailed t test (Huh-7.5 cell genome copies and DMV electron microscopy quantification) to determine statistical significance using GraphPad Prism version 9. Asterisks show statistical significance as follows: *, P ≤ 0.05; **, P ≤ 0.01; ***, P ≤ 0.001; ****, P ≤ 0.0001.

### Data Availability

The scRNA-seq data generated in this work is available at accession ID MK059949.1. Murine scRNA-seq can be accessed in NCBI BioProject with accession code PRJNA573959.

## Acknowledgments

Confocal microscopy images were acquired with the help of Dr. George Campbell at the Cell & Tissue Imaging Center which is supported by SJCRH and NCI P30 CA021765. Electron Microscopy images were acquired with the help of Dr. Cam Robinson and Nathan Kurtz using the Electron Microscopy Shared Resource (SJCRH/ALSAC and NCI P30 CA021765). These studies were supported by a NIAID grant 1R03AI166434-01 and funding from ALSAC to S.S.C., and K22 AI156116 to V.C.

---

**Fig S1.**
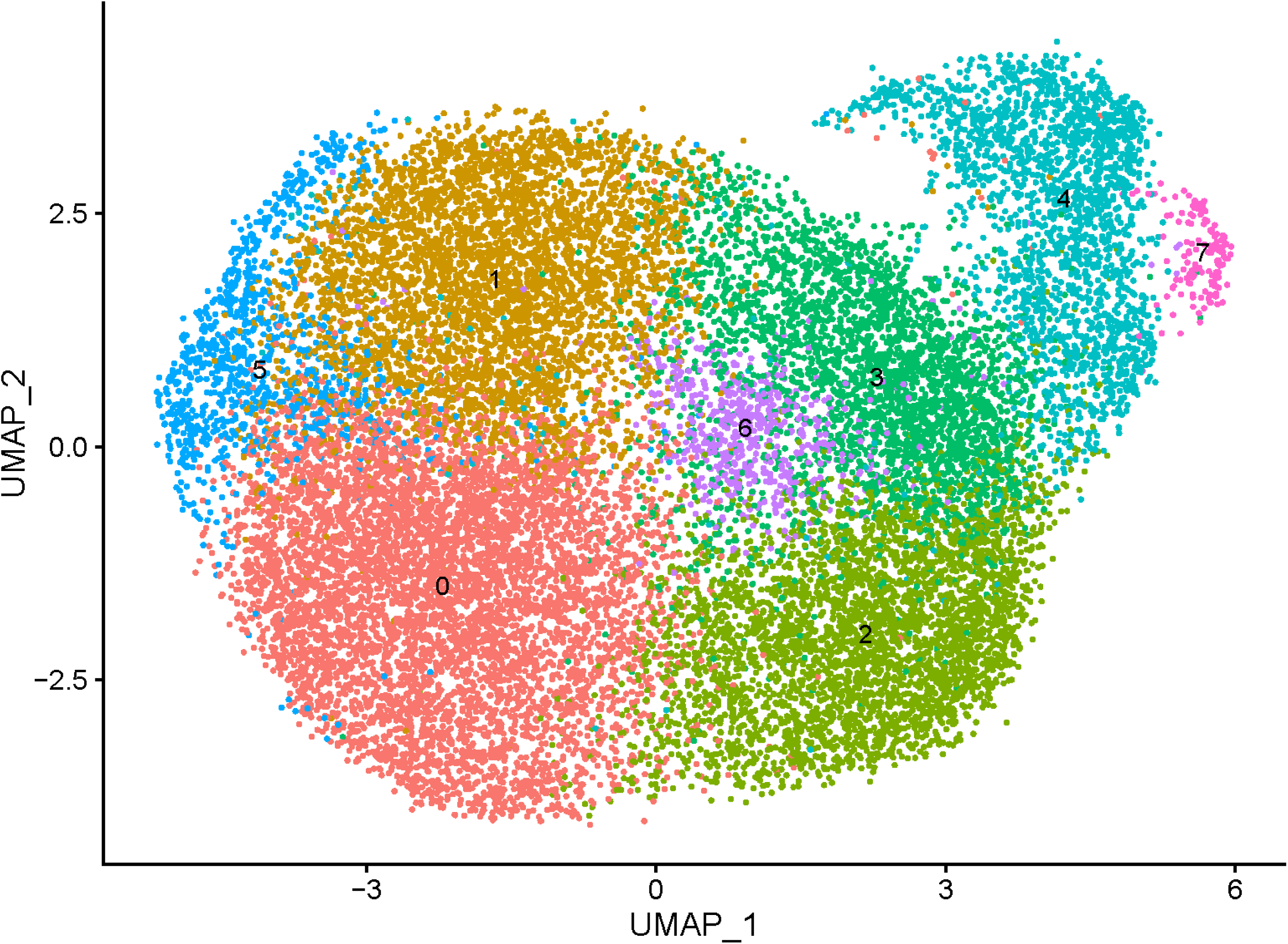
Single cell RNA sequencing UMAP. Initial clustering of 10X single cell RNA sequencing dataset shown on UMAP.

**Fig S2.**
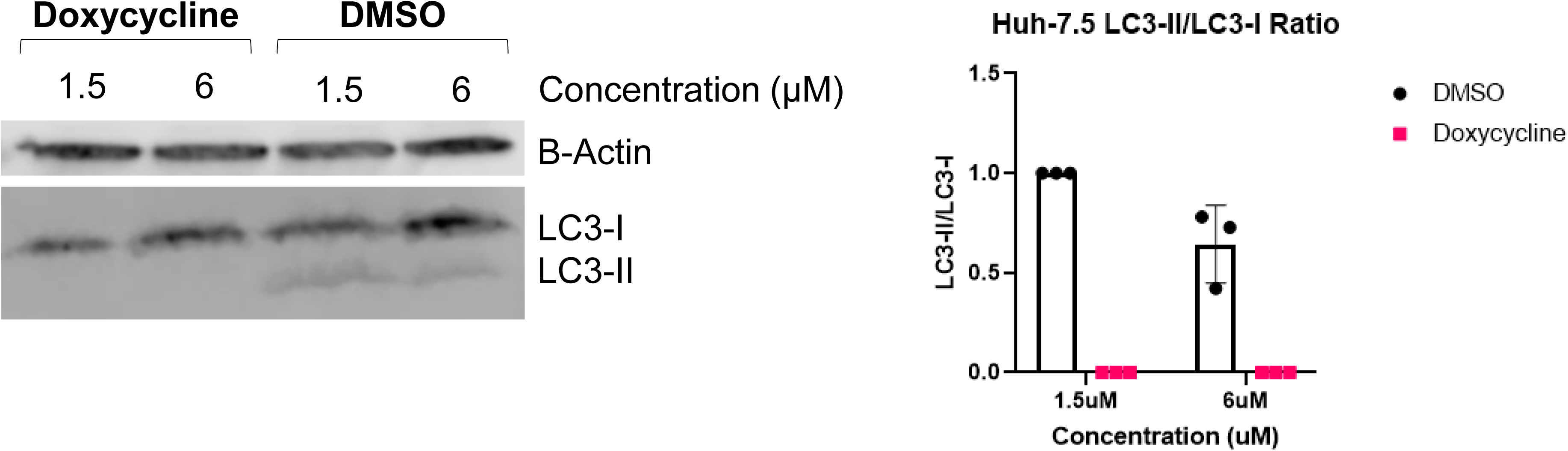
Validation of Huh-7.5 LC3 dysregulation upon doxycycline treatment. Immunoblot and quantification of LC3-II/LC3-I expression in cell lysates from Huh-7.5 cells treated with 1.5 or 6 μM doxycycline or DMSO control.

**Fig S3.**
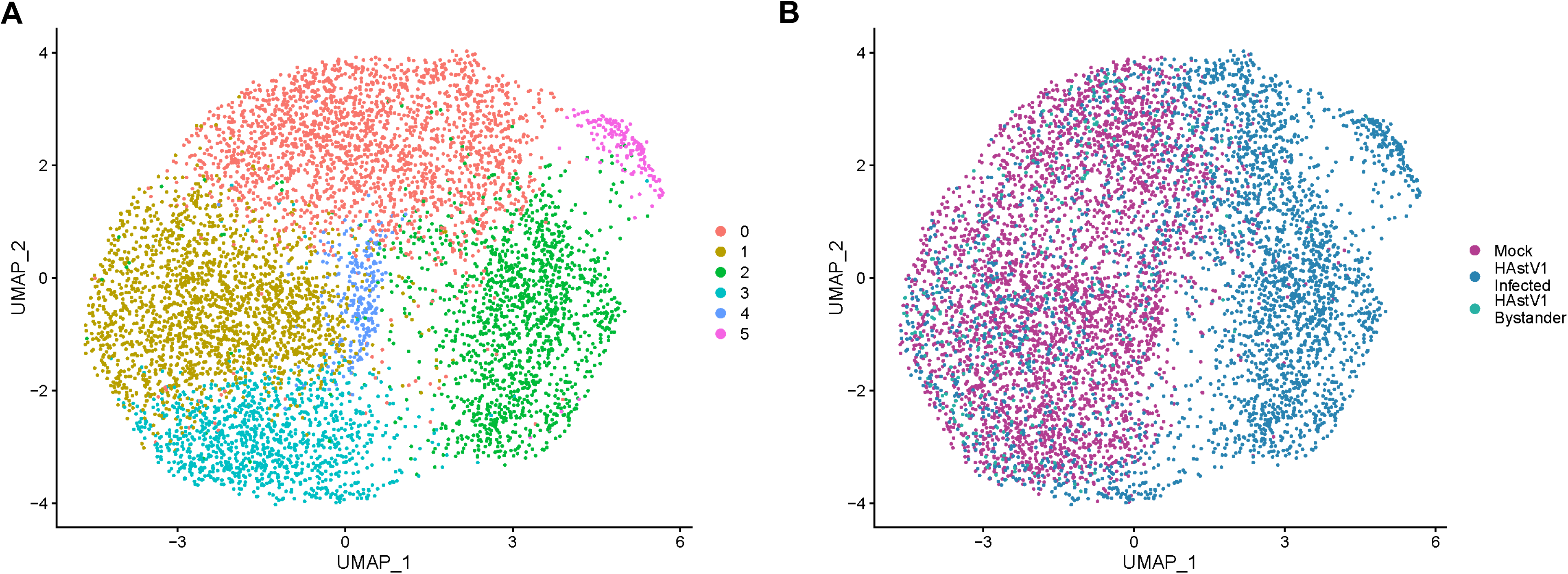
Single cell RNA sequencing UMAP for HAstV-1 and Mock 24h samples. (A) Unsupervised clustering. (B) Clustering by HAstV-1-infected, HAstV-1 bystander, and mock cells.

